# Burstprop for Learning in Spiking Neuromorphic Hardware

**DOI:** 10.1101/2023.07.25.550525

**Authors:** Mike Stuck, Richard Naud

## Abstract

The need for energy-efficient solutions in Deep Neural Network (DNN) applications has led to a growing interest in Spiking Neural Networks (SNNs) implemented in neuromorphic hardware. The Burstprop algorithm enables online and local learning in hierarchical networks, and therefore can potentially be implemented in neuromorphic hardware. This work presents an adaptation of the algorithm for training hierarchical SNNs on MNIST. Our implementation requires an order of magnitude fewer neurons than the previous ones. While Burstprop outperforms Spike-timing dependent plasticity (STDP), it falls short compared to training with backpropagation through time (BPTT). This work establishes a foundation for further improvements in the Burstprop algorithm, developing such algorithms is essential for achieving energy-efficient machine learning in neuromorphic hardware.

## 1 Introduction

The growing demand for DNN-based machine learning applications has led to increased power consumption during the training process, drawing attention to the need for more energy-efficient solutions [6]. SNNs implemented in neuromorphic hardware have emerged as a promising energy-efficient alternative to neural networks deployed in conventional computer architectures. Taking inspiration from biological neural systems, SNNs process information based on event-driven, sparse communication mechanisms to reduce computational overhead and power consumption.

There is, however, a need for effective algorithms for training SNNs in neuromorphic hardware. DNNs are predominantly trained using Back Propagation (BP)-based gradient descent. While BP-based methods can be used to train SNNs, these methods are not conducive to implementation in neuromorphic hardware, since BP requires the availability of network-wide information for gradient computation, which must be stored in memory. Rather than relying of BP, an algorithm for training SNNs in neuromorphic hardware should assign credit and coordinate plasticity based on information locally available to the synapses.

Recent theoretical work in neuroscience has proposed an approximation of BP, based on a combination of two-compartment neurons, feedback-associated burst generation and burst-dependent plasticity [5]. Together, these features implement an algorithm, Burstprop, that has been shown to enable credit assignment within hierarchical SNNs with online and stable learning. The learning task, however, has been limited to solving the XOR problem. Furthermore, this simple problem was solved using a disproportionately large number of neurons (4000 neurons in a hidden layer). The ability of Burstprop to solve more difficult problems with a smaller number of neurons remains to be shown, an essential test for any neuromorphic consideration.

In this work, we have adapted Burstprop for training hierarchical Spiking Neural Networks (SNNs) and have applied it to the MNIST digit classification task. Our implementation uses an order of magnitude fewer neurons and compares favorably with other local learning methods while remaining below the performance of offline methods. These results satisfy the necessary first step for the applicability of Burstprop to on-chip learning in SNNs.

## 2 Methods

We simulate a network of spiking neurons with a local plasticity rule, spike-timing communication and integrative properties adapted from Ref. [5] and [3].

### 2.1 Two-Compartment LIF Neurons

Each neuron consists of segregated somatic and dendritic compartments with independent membrane potentials *V*_*s*_ and *V*_*d*_, respectively. The somatic compartment follows leaky integrate-and-fire dynamics:

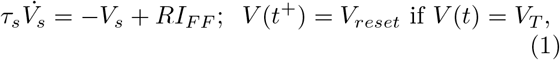

where *τ*_*s*_ = 10ms is the membrane time constant, *I*_*F F*_ is the net synaptic input current from feedforward connections, *R* = 1 is a parameter corresponding to a scaled membrane resistance, *V*_*reset*_ = 0 is the reset potential, and *V*_*T*_ = 1 is the threshold potential. The somatic compartment receives inputs from lower-level neurons, such that *I*_*F F*_ is a sum of current pulses at each pre-synaptic spike whose amplitude is sent by the synaptic weights *w*. Similarly, the dendritic compartment integrates *I*_*F B*_ leakily with the same *R* and *τ* but without a threshold, where *I*_*F B*_ performs a weighted sum of current pulses from spikes of the layer above.

The variable *V*_*d*_ does not affect *V*_*s*_ but modulates the probability *p*_*b*_ that a somatic spike is labelled as a burst, *p*_*b*_ = *σ*(*βV*_*d*_). Here *σ* is the Sigmoid function and *β* = 4 is a scaling factor added to prevent feedback saturation. Bursts will be used as credit-carrying signals transmitted from a neuron’s somatic compartment to the dendritic compartments of lower-level neurons. The algorithm uses a marked point process to distinguish between spikes and bursts and thus, does not require short term plasticity to multiplex the two signal types. Employing two-compartment neurons, along with two distinct signal types (spikes and bursts), facilitates the concurrent processing of feedforward signals and credit-carrying feedback error signals.

### 2.2 Network Architecture

The two-compartment neurons form the single hidden layer and the output layer of a hierarchical neural network. The network’s input layer transmits spike trains derived from input data to the hidden layer (Fig. 1). In the feedforward direction, layers are fully connected and undergo plasticity. In the feedback direction, the output neurons and the hidden neurons are fully connected with static randomly initialized synaptic weights. There are no feedback connections to the input neurons. All weights are randomly initialized according to a normal distribution with standard deviation proportional to the reciprocal square root of the number of presynaptic neurons. This initialization allowed the network to consistently learn effectively.

**Figure 1:**
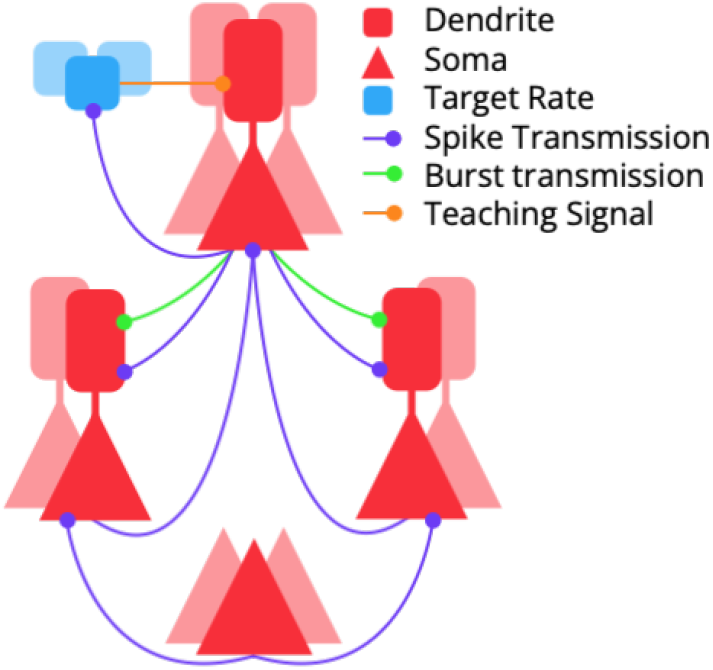
Network schematic consisting of input layer, hidden layer and output layer. Output spike rates are compared to target rates to generate a teaching signal that is sent to output neuron dendrites.

The input layer transforms data into spike trains that are transmitted to the hidden layer neurons. The output layer computes the error, which is continuously computed based on the difference between the spike rates of the output neurons and their respective target rates. This error generates a teaching signal, which is transmitted to the output neuron’s dendrites and subsequently backpropagated through the network.

### 2.3 Synaptic Plasticity Rule

Plasticity of a synaptic weight *w*_*ij*_ between a pre- and a postsynaptic neuron (labeled *i* and *j*, respectively) follow:

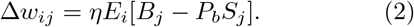

Here *η* is the learning rate, *E*_*i*_ denotes an eligibility trace of the presynaptic spikes, *B*_*j*_ represent postsynaptic bursts and *S*_*j*_ postsynaptic spikes. *P*_*b*_ = 0.5 is the baseline burst probability.

Considering that every burst is accompanied by a spike, the weight change induced by a postsynaptic spike must be multiplied by the baseline burst probability to ensure zero average synaptic plasticity in the absence of postsynaptic dendritic polarization. If the postsynaptic dendritic potential is high, the burst probability will rise above the baseline, causing the synaptic weight to increase on average. Conversely, if the dendritic potential is low, the burst probability will fall below the baseline, resulting in a decrease of the synaptic weight. The sign of synaptic plasticity is therefore controlled by the postsynaptic dendritic potential. The magnitude of synaptic plasticity depends on three factors: the presynaptic neuron activity through *E*_*i*_, the postsynaptic activity through the rate of bursts and spikes, and the postsynaptic dendritic potential that controls the burst probability. Feedback spikes are transmitted backward with intensity multiplied by the baseline burst probability to ensure an average propagating signal of zero in the absence of postsynaptic dendritic polarization.

### 2.4 Training Protocol

To train the network, each 784-pixel sample, with intensities ranging from zero to one, was converted into Poisson spike trains with a mean spike rate set to be proportional to the pixel intensity. Each sample is presented for 100 ms, during which the feedforward weights adapt according to the plasticity rule. The network included 10 output neurons, corresponding to the 10 classification categories.

During training, the output neuron corresponding to the true classification category is assigned a high target rate, while all other neurons are assigned a low target rate. This encouraged the correct output neuron to fire at a higher rate than the other output neurons. For a single training epoch, all samples from the training set were presented in a random sequence.

After each epoch, the network’s accuracy was assessed. To evaluate the accuracy, each sample from the test set was presented to the network for the same duration, and the output neuron with the highest spike count for a given sample determined the classification. Synaptic plasticity was disabled during testing to ensure the weights remained unchanged during testing. This meant that the segregated dendrite and burst signals served no purpose and did not need to be simulated. The networks were trained for a number of epochs until learning had converged.

### 2.5 Surrogate gradient optimization

For the implementation of SGD, the network architecture followed a similar structure to Burstprop, consisting of 784 input neurons, a layer of hidden neurons, and 10 output neurons. The MNIST data was presented in the same manner, with each sample represented as a 100 ms Poisson spike train. The output neurons were non-firing, and the error was calculated based on the maximum membrane potential of the output neurons during the 100 ms presentation of each sample.

During the BPTT gradient calculations, discontinuities in the forward pass were replaced with a smooth Sigmoid function to bypass the vanishing gradient issue. Each network was trained with batches of 256 samples for 100 epochs, at which point learning had converged.

### 2.6 Unsupervised STDP

For the unsupervised STDP learning method, the network consisted of a network of excitatory hidden neurons, which were connected to an equal-sized layer of inhibitory neurons. The inhibitory neurons provided lateral inhibition, serving to decorrelate the hidden neuron representation. Data, in the form of spike trains, was transmitted to the excitatory neurons and the network learned by STDP-based updates of the synaptic weights that connected these neurons. After training, excitatory neurons were assigned classes based on their activity in response to training data. In this work, the simulations of this method have not been replicated. Instead, classification accuracies are referenced from Ref. [2], where complete implementation details can be found.

## 3 Results

### 3.1 Burstprop Learning

Figure 2 depicts the learning curve for a network with 100 hidden neurons trained with Burstprop on the MNIST handwritten digit data set. The network’s performance is evaluated on a validation dataset after every training epoch, with the resulting accuracies and errors plotted accordingly. The validation error decreases and the corresponding accuracy increases with the number of training epochs until learning converges. These findings demonstrate the effectiveness of the Burstprop-trained network in successfully learning and generalizing from the MNIST dataset.

**Figure 2:**
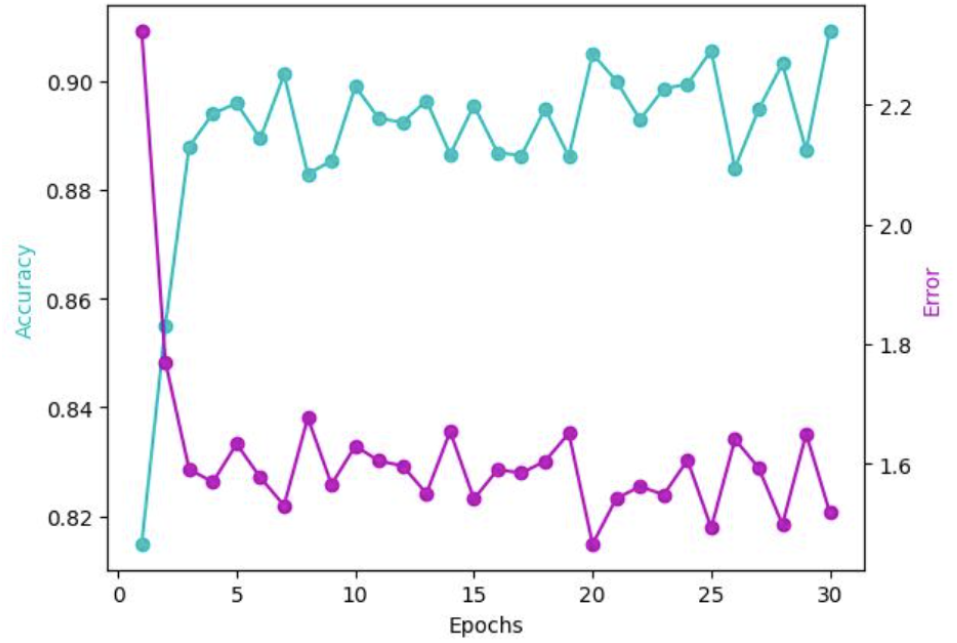
Learning curve for network of 100 neurons trained with Burstprop on the MNIST dataset. Validation error (magenta curve, right axis) and accuracy (cyan curve, left axis) are shown as a function of epochs. increases with the number of training epochs.

**Figure 3:**
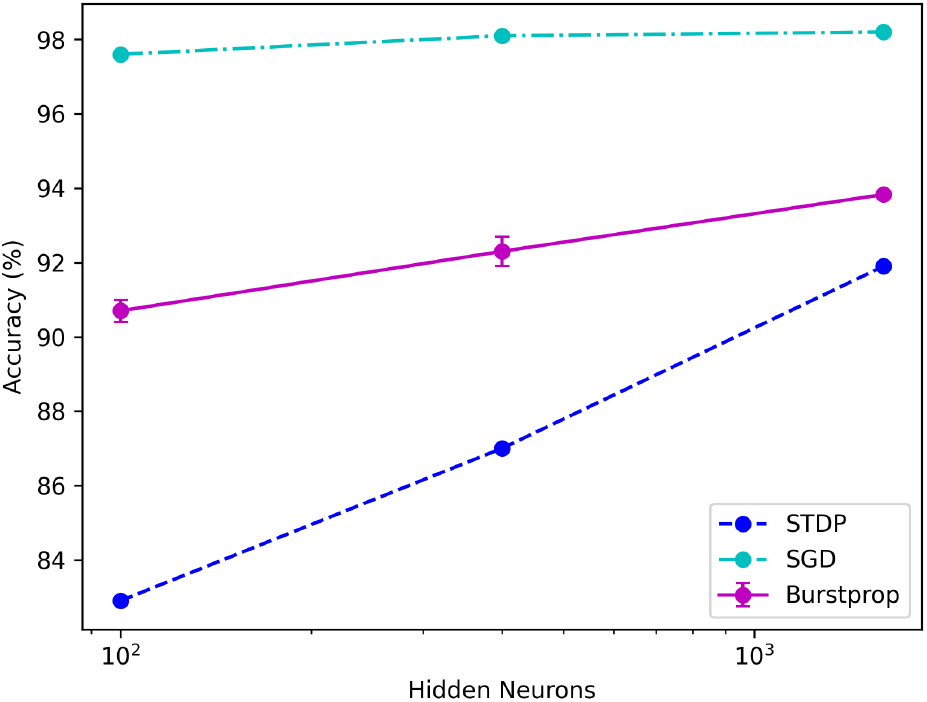
Test accuracy on the MNIST dataset as a function of number of hidden neurons for networks trained with Burstprop (magenta), SGD (cyan) and unsupervised STDP (blue).

### 3.2 Performance Comparison

To put the performance of Burstprop into perspective, we compared its performance to two other learning methods: Firstly, Surrogate Gradient Descent (SGD) with Back Propagation Through Time [4], an artificial method to train SNNs. Although this method is highly effective, it relies on the availability of network-wide information over time. This information must be stored in memory, which is not conducive to on-chip learning in neuromorphic hardware. Secondly, we compared it to an unsupervised STDP learning rule [2]. This unsupervised learning algorithm implements a local learning rule but it requires assigning a class to each neuron after training and thus relies on an additional pass over the entire dataset to track neuron activities for each sample.

The network was trained with the Burstprop algorithm for multiple trials (10 trials for 100 and 400 hidden neurons and 2 trials for 1600 hidden neurons due to resource constrainst) since the random initialization introduces some variance in performance. The average maximum test accuracy over all trials is presented on the plot. Burstprop acheives a test accuracy of 90.7%, 92.3% and 92.7% with 100, 400 and 1600 hidden neurons, respectively.

Upon comparing the classification accuracy of three distinct methods, we observed that Burstprop surpassed the performance of the STDP-based learning method, yet failed to attain a similar level of accuracy as the BPTT-based SGD method. This trend is consistent across models with 100, 400, and 1600 hidden neurons.

In all three cases, the classification accuracy improves as the number of hidden neurons increases. For the 1600 hidden neuron models, SGD achieves a classification accuracy of 98.1%, STDP attains an accuracy of 91.9. While SGD does not improve significantly form 400 to 1600 hidden neurons, Burstprop does and the trend implies that increasing the number of hidden neurons past 1600 has the potential to further increase its performance. The STDP method was also tested in Ref. [2] for 6400 excitatory neurons where it reached a classification accuracy of 95.0%. Burstprop was not tested with this number of hidden neurons due to resource constraints.

## 4 Discussion

In this work, we have furthered the preliminary tests for the use of a variation of the Burstprop algorithm to train a hierarchical network of spiking neurons for accurate image classification on the MNIST digit classification task. By multiplexing the feedforward and feedback, credit-carrying signals, synaptic plasticity can be directed based on locally available information, enabling online learning. This method provides a solution for training SNNs on-chip in neuromorphic hardware, with the goal of providing a methodology for achieving energy efficiency gains.

Although our work represents a first step in the development of an on-chip learning algorithm, a major limitation is its reduced performance compared to SGD. This presents an important avenue for future progress in improving the algorithm. Potential ideas for closing the performance gap between SGD and Burstprop include spike rate adaptation, lateral inhibition, preventing saturation of feedback signal, integration of feedback learning rules [5] and alternative error functions [4, 1].

Another aspect to consider for improving the Burstprop algorithm is its efficiency. Reducing the training time and increasing spike sparsity would in principle decrease the computational load demanded by the training algorithm, leading to reduced energy costs. These factors should be considered in any efforts to enhance the algorithm.

A significant limitation of the current implementation is the network’s single hidden layer. To learn more complex representations, a deep structure with many connected layers is necessary. Ensuring the Burstprop algorithm’s compatibility with deeper networks will require the implementation of mechanisms for avoiding saturation of feedback signal and dynamic feedback synapses to better align feedforward and feedback synapses.

## Acknowledgements

This work is supported by the Vector Scholarship in Artificial Intelligence, provided through the Vector Institute, and NSERC Discovery Grant 06872. This research was enabled by computing resources provided by the Digital Research Alliance of Canada.

